# Spatially periodic calcium profiles around single channels

**DOI:** 10.1101/077909

**Authors:** S. L. Mironov

**Affiliations:** Institute of Neuro- and Sensory Physiology, Georg-August-University, Göttingen 37073, Germany

**Keywords:** Calcium, nanodomains, periodic patterns, analytical solutions, Monte-Carlo simulations, imaging, single calcium channels

## Abstract

The concept of calcium nanodomains established around the sites of calcium entry into the cell is fundamental for mechanistic consideration of key physiological responses. It stems from linear models of calcium diffusion from single channel into the cytoplasm, but is only valid for calcium increases smaller than the concentration of calcium-binding species. Recent experiments indicate much higher calcium levels in the vicinity of channel exit that should cause buffer saturation. I here derive explicit solutions of respective non-linear reaction-diffusion problem and found dichotomous solution - for small fluxes the steady state calcium profiles have quasi-exponential form, whereas in the case of buffer saturation calcium distributions show spatial periodicity. These non-trivial and novel spatial calcium profiles are supported by Monte-Carlo simulations. Imaging of 1D- and radial distributions around single *α*-synuclein channels measured in cell-free conditions supports the theory. I suggest that periodic patterns may arise under different physiological conditions and play specific role in cell physiology.

## Introduction

Living cells are not well mixed test tubes but specialized devices where physiologically relevant events proceed in micro- or nanocompartments. This allows to efficiently isolate complex biochemical cascades from bulk interior, saving the time, space and reagents. For example, the synaptic transmission, secretion, contraction etc. utilize fast, big and local single calcium transients that are spatially limited by putative calcium-binding proteins (Augustine et al. 2003; Eggermann et al. 2011). The compartmentalization is well suited to selectively activate low-affinity calcium sensors that are often strategically positioned in the immediate vicinity of the calcium channels. The existence of highly localized calcium increases around single channels has been predicted by Neher (1986). He treated calcium binding to cytoplasmic buffers as fast and irreversible reaction without buffer saturation and considered steady state calcium profiles. Under these assumptions the reaction-diffusion (RD) problem reduces to the linear ordinary differential equation (ODE)

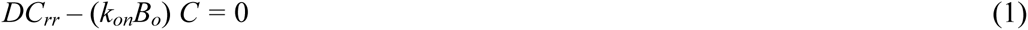

where *C*= [Ca]/*r*, *C*_*rr*_ is the second derivative by the radial coordinate *r*, *D* = 220 μm^2^/s is calcium diffusion coefficient in the cytoplasm, *B*_*o*_≈ 0.2 mM is the total concentration of cytoplasmic buffer(s) and *k*_*on*_ ≈ 2.10^8^ M^−1^s^−1^ is the on-rate constant for calcium binding (Mironova & Mironov, 2008). The solution of (1) is straightforward

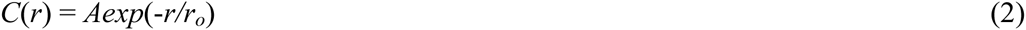
and predicts exponential decay of calcium levels from the point source, a single calcium channel. A typical space constant is 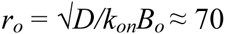 and the characteristic time-constant is *τ*_*o*_ = 1/*B*_*o*_*k*_*on*_ = 40 µs, much lower than typical open and closure times of the calcium channels (>0.1 ms). Thus steady state profiles around single channels should be established fast during opening and quickly disappear during closures. The mean time for calcium unbinding from the buffer is *τ*_*off*_ = 10 ms and can be omitted from the consideration.

The problem yet appears when we consider maximal calcium levels around the channels. For radial diffusion of calcium it is defined as

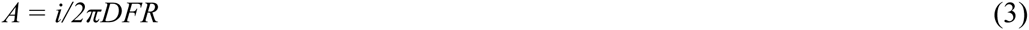
where *i* is the single channel current *i* and *R* is exit radius of the channel. A theoretical estimate of the ratio *A/i* is 1.2 mM/pA (Mironov, 1990) and close to recent experimental values ≈0.7 (Tay et al. 2012) and 1.0 mM/pA (Tadross et al. 2013). Therefore local calcium increases can be really big that does not validate the use of a simple linear model. Indeed, for single calcium currents *i* > 0.2 pA, the calcium level at channel lumen exceed a typical concentration of cytoplasmic buffers (0.2 mM) and violates the assumptions adapted in linear treatment.

The considerations prompted me to revisit the problem of steady state calcium profiles around single calcium channels. A general problem of calcium diffusion into the cytoplasm with multiple calcium binding proteins (buffers) is described by the system of nonlinear parabolic partial differential equations (PDE)

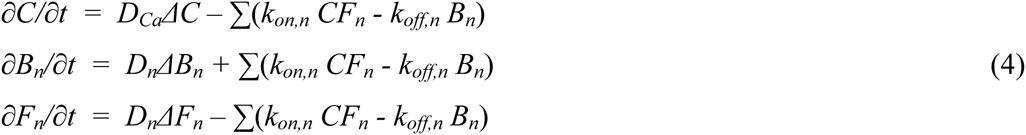
where *∆* stands for the Laplacian; *C*, *F*_*n*_ and *B*_*n*_ are the concentrations of calcium and *n*th buffer in free and calcium-bound forms, respectively; *k*_*on,n*_ and *k*_*off,n*_ are the rate constants of calcium binding to and dissociation from the buffers. The ratio *k*_*off,n/*_*k*_*on,n*_ = *K*_*d,n*_ is the dissociation constant of the buffer and measures its affinity to calcium. *D*_*Ca*_ is the calcium diffusion coefficient and *D*_*n*_‘s are the diffusion coefficient of the buffers.

To make the problem tractable, I first consider a one-dimensional steady state problem of calcium diffusion in the presence of single buffer and derive explicit solutions without any restrictions on the magnitude of calcium influx or buffer saturation. All derivations below are made in non-dimensional form. The concentrations are normalized to the total buffer concentration *B*_*o*_ (0.2 mM), and the times and distances are defined as *t* = *t/τ*_*o*_ and *x* = *x/r*_*o*_ (see the definition of the characteristic scales above). For calcium and single buffer the system (4) is reduced to

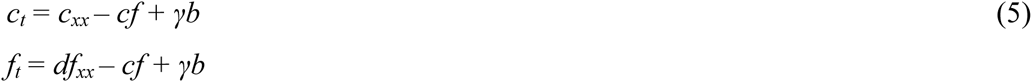
where *c* and *f* are the concentrations of free calcium and buffer, respectively, and *d* = *D*_*B*_/*D*_*Ca*_ is buffer diffusion coefficient relative to that of calcium. The last term in the right-hand side represents calcium unbinding from the buffer. The value *γ* = *k*_*off/*_*k*_*on*_*B*_*o*_ = 0.005 is very small and can be neglected. Subtraction of the two equations in (5) gives a single PDE

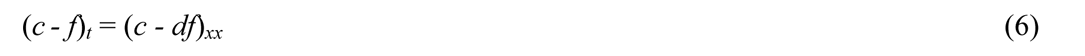
that is readily solved in the case *d* = 1 (*D*_*B*_ = *D*_*Ca*_) with

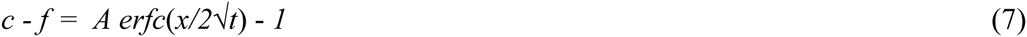
where *erfc*(*y*) is the complementary error function. The constant *A* is defined by the boundary conditions implying that both calcium and buffer at the channel lumen are free (not yet reacted).

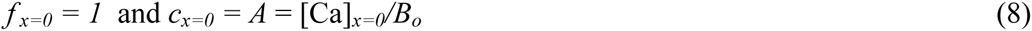

Constant calcium at channel lumen in 1D-presentation is equivalent to the assumption of constant calcium flux through the channel in the case of radial diffusion (Neher, 1986; Mironov, 1990). The radial calcium gradients in the linearized treatment can be obtained by dividing 1D-spatial profiles by the distance from the channel (Mironov, 1990). In a non-linear case this transformation is not exact but still provides a very good approximation as shown in the Appendix. Expressing *f* through *c* from (7) transforms the first Eq. (5) into a single PDE

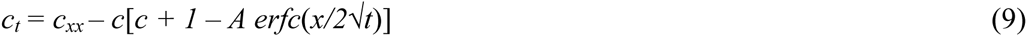

More compact presentation is obtained after scaling the variables as

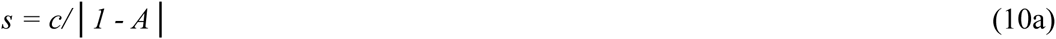

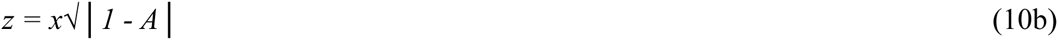

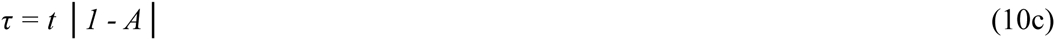
that transforms (9) into

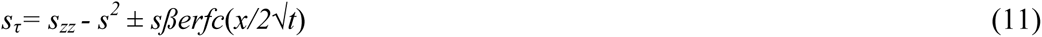
with *β*=*A*/|*1* – *A*|. Note that from a single PDE we obtained two equations. This is due to the fact that normalisation factor (*1 – A*) can be either positive or negative. For *A* > *1* the square root in (10b) without modulus is imaginary. Introduction of modulus is equivalent to introduce (±) sign before the linear term *sβerfc.* This has a crucial importance because (11) possesses the two different solutions that describe spatial calcium profiles below and above the critical point *A = 1*.

PDE (11) is non-linear and its general solution is yet to be found. It is possible to obtain approximate analytical solutions as shown below. I start first with the analysis of steady state problem that already gives important insights. By approach to the steady state *s*_*t*_ → 0 and *erfc →1* and (11) is replaced by the ODE

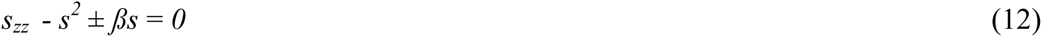
that is solved by subsequent integration. The first one gives

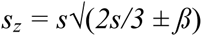
and the results of second integration are

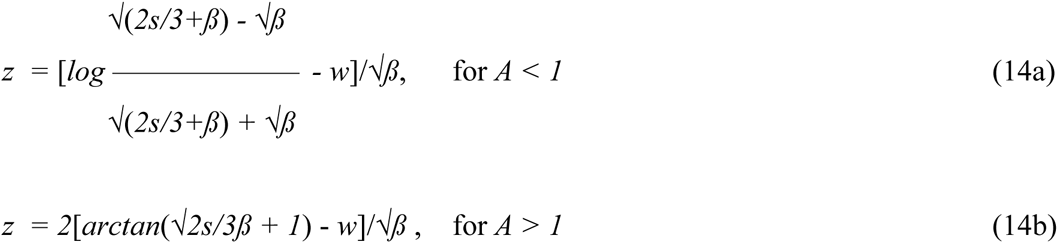

Here *w* is the constant of integration determined by the boundary condition. Eqs. (14) define the concentration of calcium implicitly. Inversion of (14) gives explicit expressions

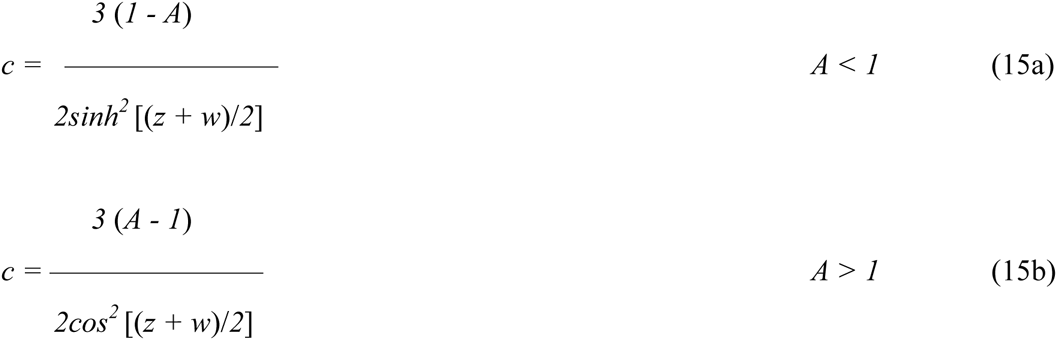

Note that the concentrations and distances are given in the normalized variables 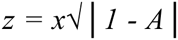 and *c* = [*Ca*]/*B*_*o*_. The constant of integration *w* has a meaning of phase shift.

As mentioned above, the appearance of the two solutions stems from inherent structure of stationary RD equation (12). The result is general and is not due to the boundary conditions imposed or assumption about equal diffusion coefficients. Let the relative diffusion coefficient for buffer be *d* = *D*_*B*_/*D*_*Ca*_ < 1. In the steady state Eq. (6) is

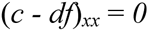
and prescribes (*c — df*) = *const* = *A – 1*, by analogy to (7). Insertion of *f* = (*c — A + 1*)*/d* into the first Eq. (5) gives the steady state equation, identical to (12). Therefore concentration dependences are still given by (15), given a new scaling

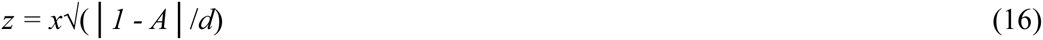

The characteristic scale constant is now 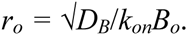. In comparison with that defined in (2) here *D*_*Ca*_ is replaced with *D*_*B*_. Thus, for *D*_*B*_ < *D*_*Ca*_ the width of calcium nanodomains depends from the diffusion coefficient of the buffer, not calcium. The difference can be significant for cytoplasmic calcium buffers that are bulky proteins. For calmodulin and calbindin *D*_*B*_ = 0.03*D*_*Ca*_ (Mironova & Mironov, 2008) and the characteristic space scale should increase by around 6-fold.

It is important to know, how the predicted steady state calcium profiles develop. As PDE (11) is not yet solvable, I examined its asymptotics by setting *s*_*t*_ = *0*, an assumption used sometimes in the RD field. This leads to the non-linear Schrödinger equation

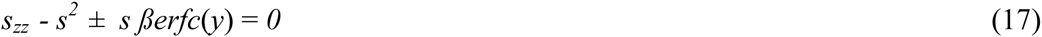
where 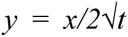 is a classical Boltzmann similarity variable that naturally appears in various diffusion problems (Polyanin & Zajtsev, 2007). Without quadratic term, Eq. (17) may be considered as typical problem of quantum mechanics describing the particle moving in the potential *βerfc*(*y*). No analytical solution exists, but WKB approximation (Holmes, 1995) delivers a leading term in in solution

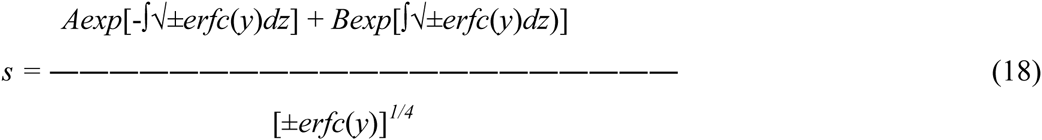

This analytical result is important, because it also predicts the two types of solutions - decaying and periodical. They appear because the expressions under the square root in (18) are either positive or negative that give either real or imaginary arguments in the exponentials. At big times, when *t→∞*, the function *erfc*(*y*)*→ 1*, and 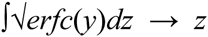 and 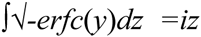. Therefore the sum in the numerator invokes either hyperbolic or trigonometric functions.

This analysis is confirmed by numerical solution of (17) using the shooting method (Polyanin & Zajtsev, 2007). First, the calculations showed that *s*_*t*_ << *s*_*zz*_ at times *t* > 0.01 ms. This validates use of asymptotic approach to derive (17). The time-dependent solutions converged to the steady state profiles given by (15). Fig. 1A shows the development of calcium profiles for decaying and periodic solutions. For *A* = 0.2 the exponential profile is established within < 1 ms. For *A* = 2 the initial decaying pattern transforms into a spatially periodic waveform. The same approach was used to calculate radial calcium profiles (Fig. 1B).

**Fig. 1.**
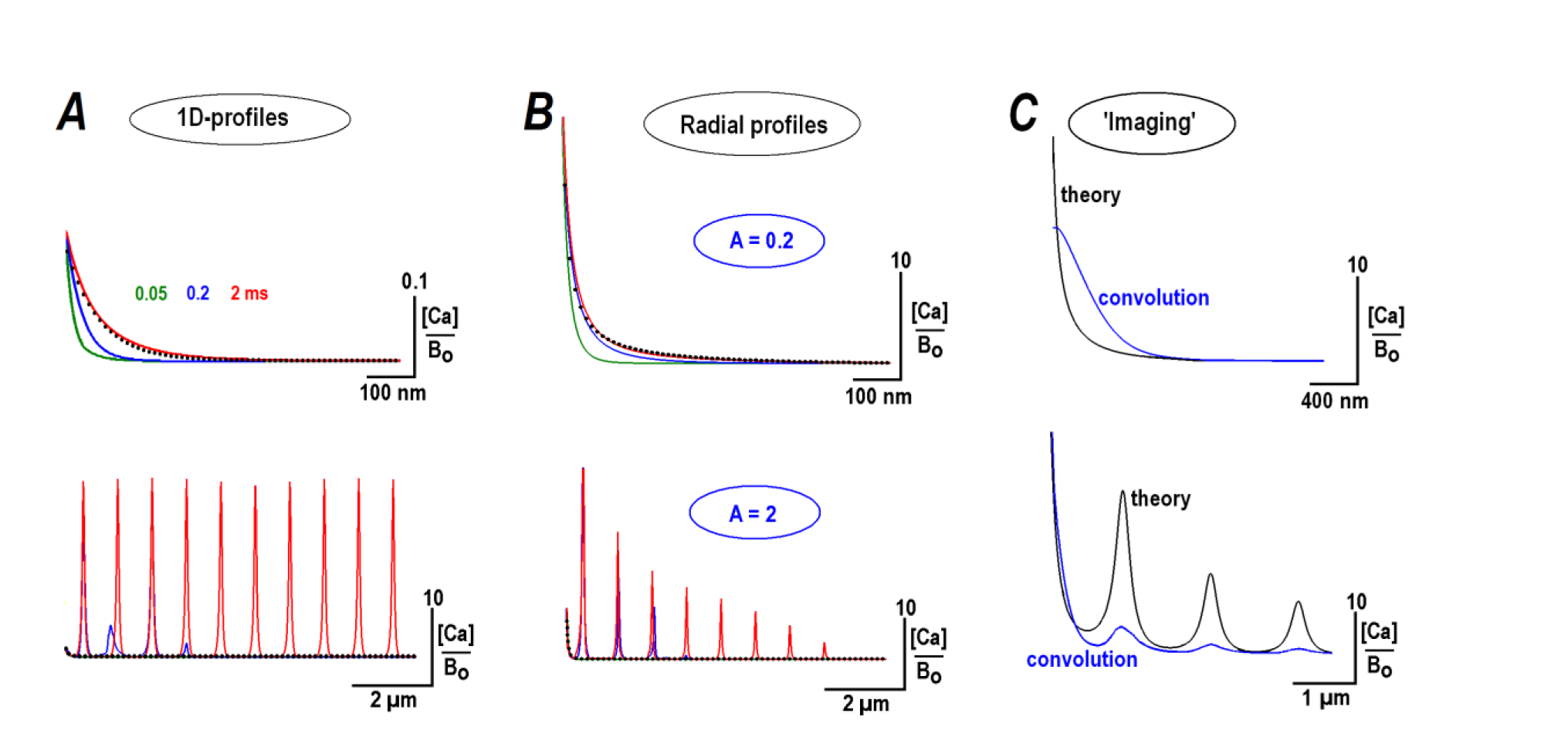
Steady state profiles for calcium diffusing from the point source. One-dimensional (***A***) and radial (***B***) calcium distributions around the point source (single calcium channel) obtained as the solutions of non-linear RD equation (12). Relative calcium concentrations at the point source [Ca]_*o*_/*B*_*o*_ bracket the critical value *A = 1* that separates decaying and periodic solutions. The time-dependence was obtained by solving Eq. (17) and the upper and lower graphs show calcium profiles at 0.05, 0.2, and 2 ms as curves with respective colours. The traces at 2 ms coincide with the steady state distributions given by Eqs. (12). The black dots indicate stationary exponential solutions predicted by linearized treatment, Eq. (1). ***C***- To simulate blurring of radial calcium profiles by imaging, the theoretical curves (black) were convoluted with the Gaussian point spread function (HWHM = 0.4 µm). Note that despite blurring, the secondary peaks for *A* = 2 are clearly visible.

How would these theoretical calcium profiles look in experiments? Calcium profiles measured with indicator dyes are inevitably distorted due to finite resolution in imaging. To simulate such effects, I convolved the radial calcium profiles with the Gaussian point spread function 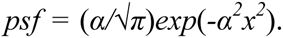. The half-width-half-maximum (HWHM) was set to 0.4 μm, close to the experimental resolution of imaging system used to measure calcium distributions around single channels (see below). As expected, the ‘imaging’ blurs calcium distributions, the amplitudes of radial calcium gradients decrease and they broaden (Fig. 1C). For decaying transients (*A < 1*) the width of calcium nanodomains is around HWHM. Periodic radial patterns for (*A > 1*) are also deformed but the secondary peaks are well discerned.

I next simulated stochastic diffusion of calcium in the presence of buffer (see Methods). Calcium ions appeared randomly at the origin (*x* or *r = 0*) and the mean level was set to a prescribed *A* value. Representative simulation runs are presented in Fig. 2 as kymographs showing instantaneous positions of particles (free calcium, free and bound buffer). Initial and final distributions of species were obtained as averages of 1000 subsequent ‘frames’ in the beginning and the end of the runs. In all simulations the distribution of calcium in the beginning was always decaying and did not appreciably change with time for *A* = 0.2 (Fig. 2A). A half-width in this case was *r*_*o*_ ≈ 0.1 µm, close to the expected value. For *A* = 2, a secondary peak around 0.6 µm slowly evolved (Fig. 2B), in accord with a theoretical estimate under these conditions.

**Fig. 2.**
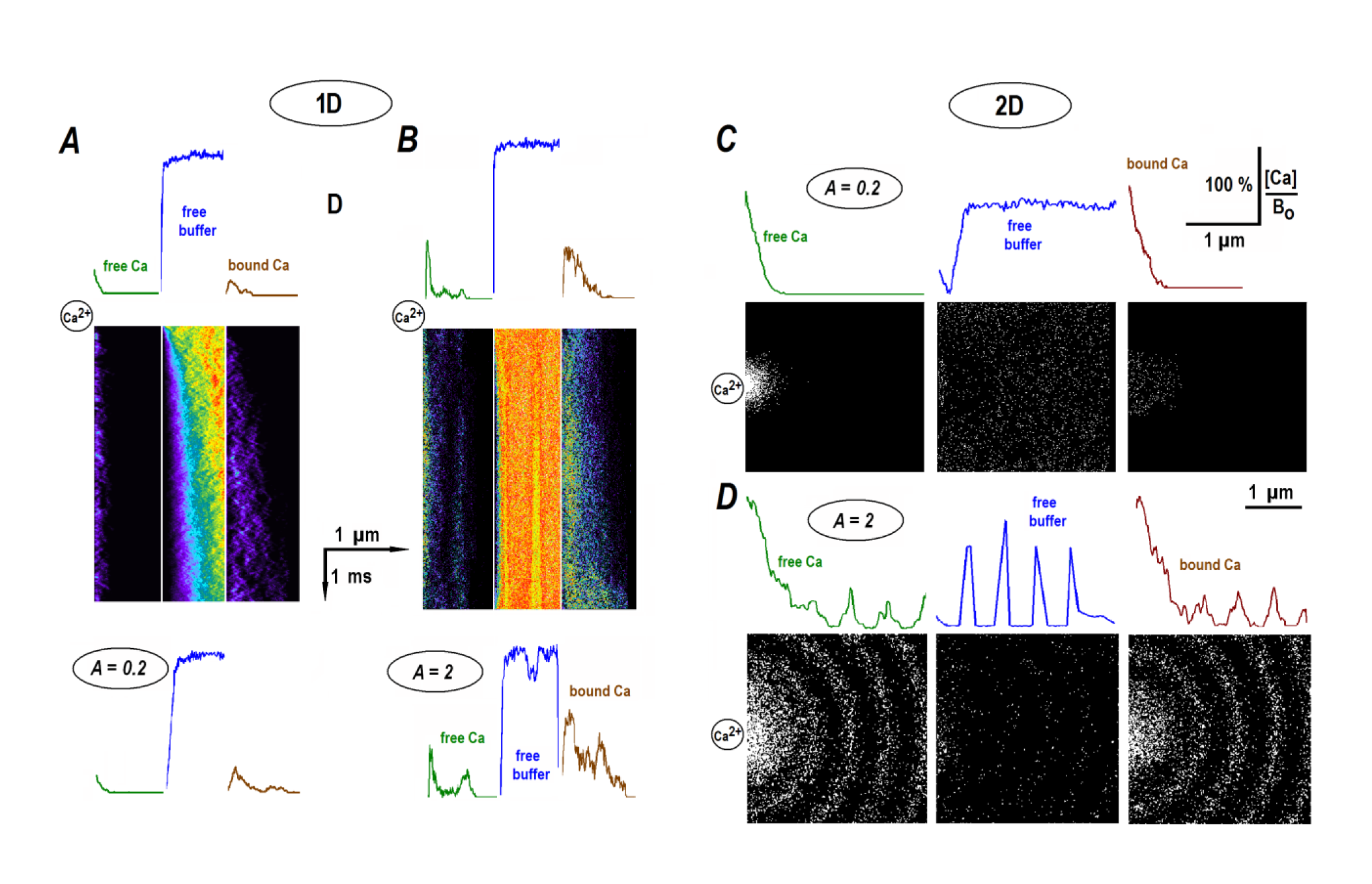
Stochastic simulations of calcium distributions in the presence of buffer. Simulations were performed as described in Methods. ***A***, ***B —*** The kymographs plot 1D- instantaneous distributions of free calcium and buffer species. The time and space directions are indicated by the arrows in the middle. Calcium ions appeared randomly at the origin (the upper left corner in the kymograph), spread and reacted with buffer molecules. The representative runs were performed at different ratios *A* = [Ca]_o_/*B*_*o*_ = 0.2 (***A***) and 2 (***B***). The concentration profiles above and below the kymographs plot averaged initial and final distributions of particles in the runs. ***C***, ***D*** — Radial profiles for 2D-diffusion. The snapshots in panels were taken in the beginning of the run and after 1 ms. Calcium influx was set constant and the ratio between calcium and free buffer was set to *A* = 0.2 and 2. The curves above the panels present radially averaged ‘steady state’ distributions of free calcium and buffer as well as bound calcium.

In order to test theoretical predictions, I imaged calcium profiles around single α-synuclein (αS) channels in the excised patches. A cell-free experimental configuration is better suited to imaging calcium nanodomains on several reasons. First, the composition of bath solutions is well defined. Second, I used fluo-4 that has weak intrinsic fluorescence in a free form and increases it >10-fold after calcium binding. Therefore the regions where calcium binding to the indicator does not occur, marginally contribute to measured fluorescence, minimize out-of-focus effects and improve spatial resolution. The on-cell measurements, in contrast, can be severely contaminated by bulk fluorescence that considerably masks local changes in calcium.

αS channels were incorporated into the membrane of hippocampal neurons as described (Mironov, 2015) and calcium distributions were measured with fluo-4 using TIRF excitation (Methods). Mean calcium levels from inside-out patches depicted well opening of αS channels (Fig. 3, *top panels*). αS-channels have three conductance states, but the upper two are short living and did not significantly contribute to calcium changes. Steady state distributions were imaged at different holding potentials set to desired values of the theoretical parameter *A* in Eq. (8), the ratio between calcium levels at channel lumen and buffer concentration in the medium (100 µM fluo–4). Under conditions used, *A* values are numerically equal to the mean single channel current e. g. for *i* = 0.1 pA, *A* = 0.1 etc.

**Fig. 3.**
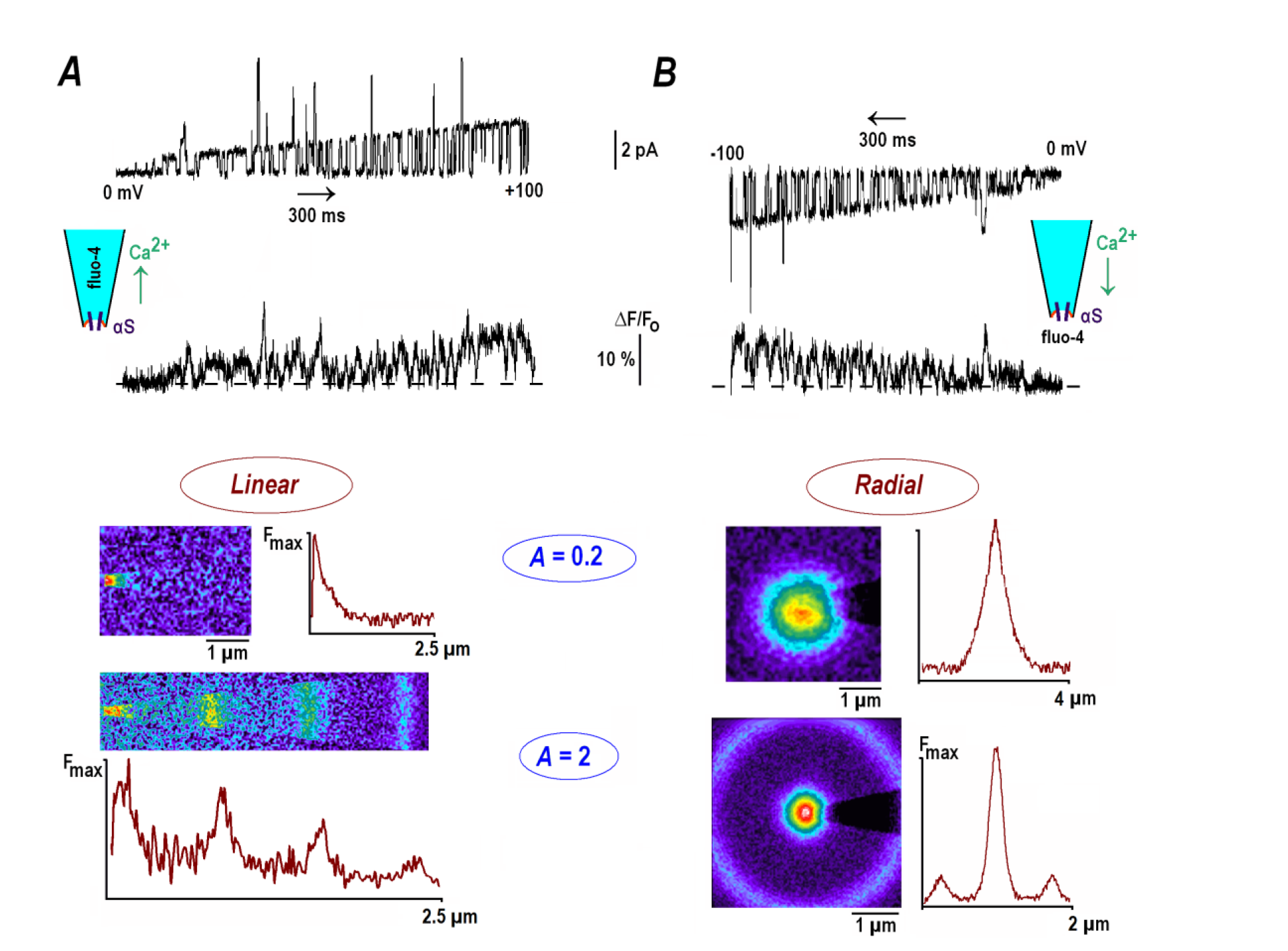
Calcium gradients established around α-synuclein channels in inside-out patches. The channels were incorporated into the patches excised from cultured hippocampal neurons as described in Methods (see also Mironov, 2015). ***A*** — The outward calcium currents through αS channels, mean calcium changes and 1D-profiles. The uppermost trace presents channel opening during 2 s-voltage ramp from 0 to +100 mV. The trace below shows relative changes in mean fluo- 4 fluorescence in 2 µm-circle around the patch. Pipette solution contained 100 µM fluo-4 in 154 mM NaCl and 88 mM CaCl_2_ was present in the bath. Fluo-4 images (acquisition time, 3 s) indicate stationary calcium distributions at different patch potentials. The parameter *A* is the ratio of calcium level at channel lumen (determined by the single channel current, Eq. (3)) and buffer concentration. Horizontal linescans present stationary 1D-calcium distributions. For *A* = 0.2 (mean single channel current, 0.2 pA), HWHM (half-width at half-maximum) was 0.43 µm, close to the radial resolution of the experimental set-up (HWHM ≈ 0.4 µm, Methods). For *A* = 2 the calcium distribution is spatially periodic and equidistant peaks are separated by 0.66 µm, in accord with a theoretical estimate 2*πr*_*o*_ = 0.67 µm. ***B*** - The inward calcium currents through αS channels, mean calcium changes and radial calcium profiles. The top trace shows channel activity during 2 s-voltage ramp from 0 to –100 mV and the lower trace presents mean changes in fluo-4 fluorescence around pipette tip. The fluorescence images in panels were acquired at different patch potentials and the traces present their diagonal linescans. The half-width of the spot for *A* = 0.2 was 0.41 µm. For *A* = 2, calcium distribution showed also a secondary peak. A concentric shell has radius 0.68 µm.

The one-dimensional profiles were generated by forcing calcium to diffuse from the bath into the pipette filled with fluo-4. Stationary calcium increases established fast and were localised to the pipette tip or formed periodic patterns within it that depended on the value of *A* fixed (Fig. 3A). The radial patterns generated by calcium diffused out of the pipette into the bath with fluo-4 showed single spot and had a concentric shell around it at *A > 1* (Fig. 3B). For small calcium currents (*A* = 0.2), the half-width at half-maximum (HWHM) was 0.42 ± 0.05 (linear diffusion) and 0.44 ± 0.06 µm (radial diffusion), respectively (mean data from four patches in each case). The values are close to the radial resolution of the experimental set-up (HWHM = 0.39 ± 0.03 μm, see Methods) indicating that a true quasi-exponential calcium decay is apparently hidden within the optical spot (a theoretical characteristic space constant in this case is 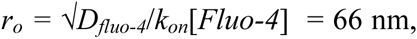 Eq. (16)). This is in line with the results of ‘imaging’ simulations (Fig. 1C). For big calcium fluxes (*A* = 2), the steady state patterns were periodic. In linear case the mean interval between the equidistant peaks was 0.65 ± 0.03 µm (*n* = 4). For radial diffusion the distance between the main and secondary peak was 0.68 ± 0.04 µm (*n* = 4). Both values are close to the theoretical estimate 2*πr*_*o*_ = 0.66 µm calculated for experimental conditions used.

## Discussion

Neher (1986) predicted existence of local gradients (calcium nanodomains) around single channels and this concept emerged into a cornerstone of contemporary physiology. It is frequently used to explain the mechanisms of key physiological processes such as secretion, synaptic transmission and muscle contraction. To describe distributions of free calcium around single channels, he used a linear reaction-diffusion model. Recent experiments (Tay et al. 2012; Tadross et al. 2013) however indicate calcium levels around 1 mM near the channel lumen. Such values exceed concentration of putative cytoplasmic buffers (~0.2 mM).

This prompted me to make a leap beyond linear model and I solved a non-linear reaction-diffusion problem for steady calcium state profiles. The solutions indicated more complex patterns of calcium distribution around single channels. The main theoretical result in Fig. 1 shows that profiles haveeither decaying to periodic forms, whose appearance depends on critical parameter *A* = [Ca]_*o*_/*B*_*o*_, the ratio between the calcium level at channel lumen and total buffer concentration. For *A < 1* calcium decays quasi-exponentially and resembles a classical solution of linear model. For *A > 1* the non-linear RD model predicts spatially periodic calcium profiles. This is a novel, unexpected and perhaps counterintuitive finding. It would be logical to assume that calcium distributions always have the same waveform and are only proportionally scaled according to the magnitude of calcium flux. A theoretical analysis does not support the mechanistic interpretation, however. When calcium levels at channel lumen exceed buffer concentration, the spatially periodic steady-state patterns appear.

Mechanistically, the existence of the two distinctly different patterns stems from the fact that the two different equations describe the profiles in the case of small and big calcium fluxes. This is instantly seen from Eq. (12). Without a quadratic term a respective ODE *s*_*xx*_ ± *s* = *0* has different sign of linear term. When it is negative (*A < 1*), the solution is exponential and when it is positive (*A > 1*), the solution is trigonometric. The result is general, but not as exotic as it may seem. A spatial periodicity is well-known in the reaction-diffusion field (Koch & Meinhardt, 1994; Vanag & Epstein, 2007) with a seminal example presented by Liesegang periodic patterns (1896) that can be produced even within a test tube.

Visualization of spatial calcium distributions in living cells at nanoscale is difficult due to considerable image distortions (Fig. 1C). Measured single calcium transients yet frequently show the main peak with broad shoulders (Beaumont et al. 2005; Shuai & Parker, 2005; Demuro & Parker, 2006) that may hide secondary calcium peaks predicted by a non-linear model. I imaged calcium profiles established around single αS-channels (Fig. 3) in cell-free conditions. The experimental configuration enabled to depict both one-dimensional and radial periodic calcium profiles. Theoretical analysis is further supported by the Monte-Carlo simulations (Fig. 2).

What could be possible implications to the cell biology? The ratio *A* =[Ca]_*o*_/*B*_*o*_ is critical for appearance of spatially periodic calcium profiles. Recent experimental estimates give for the ratio between calcium level at channel exit and the single channel conductance (*i*) the values [Ca]_*o*_/i ~ 1 mM/pA (Tay et al. 2012; Tadross et al. 2013). For *B*_*o*_ = 0.2 mM the critical value *A* = 1 corresponds to *i* = 0.2 pA. The borderline can be easily crossed when the single channel current gets bigger or channel exit radius gets smaller. Another way to increase the ratio [Ca]_*o*_/*B*_*o*_ is to take into account the effects caused by surface charges near the channels (Kostyuk et al. 1982). Even for a very moderate local surface potential –25 mV, the calcium levels under membrane are *e*^2^ ≈ 7 higher and for tetra-anion e. g. fluo-4 are *e*^4^ ≈ 50—fold lower than in the bulk that will increase the ratio *A* = [Ca]_*o*_/*B*_*o*_ dramatically. The possibilities show only a few ways to increase *A* that will promote the appearance of spatial periodic profiles around single calcium channels.

According to (16), a decrease in buffer diffusion coefficient increases spatial scaling. Calcium gradients therefore may be more extended in the presence of genetically encoded calcium probes or intrinsic cytoplasmic buffers, the bulky proteins that have much smaller diffusion coefficients. Organic calcium buffers and indicator probes have bigger diffusion coefficients (yet smaller than that of calcium). Their presence in the cytoplasm should restrict calcium increases in comparison with that established in native environment. Therefore imaging with synthetic dyes likely overestimates the width of calcium profiles.

The concept of nanodomains has been originally invoked to explain the difference in the effects of calcium buffers - EGTA and BAPTA - upon calcium-activated K^+^ channels. The two species were accordingly dubbed as slow and fast buffers, because they bind calcium with apparently 100-fold difference in the on-rate constants (Smith et al. 1984). A seemingly slow calcium binding by EGTA has been previously discussed (Mironova & Mironov, 2008). Briefly, a doubly protonated EGTA (H_2_EGTA^2−^) is dominant at physiological pH but it cannot bind calcium efficiently — a *K*_*d*_ = 4 M (Smith et al. 1984) indicates its extremely low affinity to calcium. HEGTA^3-^ species (*K*_*d*_ = 5 µM) can do this, but in 10 mM EGTA their concentration is only 0.1 mM. At ms-time scale calcium ion is captured by the first buffer molecule it meets in the cytoplasm with the on-rate close to the diffusion limit. The rate of calcium binding is *k*_*on*_[Buffer] and for equal nominal EGTA and BAPTA concentrations, an apparent 100-fold difference in *k*_*on*_ values simply reflects a 100-fold smaller concentration of calcium-binding species in the former case. At bigger times EGTA accommodates calcium in exchange for protons that are well buffered. The process is much slower and explains why EGTA should minimally disturb fast calcium transients but prevent the cell from calcium overload in e. g. whole-cell recordings, where EGTA virtually eliminates possible deleterous long-lasting increases in cytoplasmic calcium.

The predicted periodic solutions do not violate the concept of calcium nanodomains as large peaks around the channel exit are also present for *A* > 1. There are yet secondary maxima at multiples of 2*πr*_*o*_ ≈ 1 µm. They are better visible in 1D-systems, where they have a constant height. What physiological function in the living cells they may have? For radial diffusion the amplitudes of secondary peaks decay hyperbolically (~*1/r*, Fig. 1), but they are yet big enough to trigger specific calcium-dependent events in the neighbourhood. The two possibilities are worth to consider. First, a collective activity of calcium release channels in internal stores produces such events as sparks, puffs, etc. (Wang et al. 2004). They are generated by closely apposed IP_3_ receptors (=calcium release channels) that require calcium for a full activation. It is plausible that secondary calcium peaks generated by one channel will promote the activation of its neighbours. Asynchronous transmitter release (Kaeser & Regehr 2014) may represent another implication. This type of release is triggered by brief spontaneous calcium channel opening at resting potentials. The events have short duration (0.1 ms), but single channel currents are big enough to produce secondary calcium peaks that can trigger exocytosis of vesicles loosely coupled to calcium channels.

Taken together, this study reveals novel unexpected profiles that can be established around single calcium channels. The results may be fruitful in exploring new research directions and help to explain the features previously interpreted invoking more sophisticated proposals.

## Methods

### Patch-clamp and imaging

Cultures of hippocampal neurons were prepared from 2− to 4-day-old mice as described previously (Mironov, 1995). Bath and pipette solutions contained 30 mM Tris buffer (pH 7.4), and 154 mM NaCl or 88 mM CaCl_2_. The solutions had osmolality from 305 to 315 mosmol/l. Fluo-4 was from Invitrogen (Darmstadt, Germany). Patch-clamp pipettes had long shank (around 5 mm) and 20 ± 3 MOhm resistance. Imaging was made with 63x objective lens (N. A. 1.4) of an upright microscope (Axioscope 2, Zeiss). The fluorescence was excited by 488 nm light from SLM Diodenlaser (Soliton, Gilching, Germany) and captured by cooled CCD camera (BFI Optilas, Puchheim) operated under ANDOR software (500 x 500 pixels at 12 bit resolution). The laser beam was delivered from below at the angle appropriate to evoke TIRF. The spatial resolution of the experimental set-up was estimated by imaging fluorescent beads (40 nm diameter). Their half-width at half-maximum (HWHM) was 0.39 ± 0.03 µm (a mean from 12 objects).

Coverslips with hippocampal neurons were placed on the microscope stage. The inside-out patches were excised and, when they showed no activity of intrinsic ion channels, αS channels were incorporated into membrane as described previously (Mironov, 2015). The pipette was positioned nearly horizontal and carefully lowered down to the bottom. The approach was controlled by monitoring resistance, similar to that used in the scanning ion conductance microscopy (Kornchev et al. 2000). When the resistance dropped by 1%, it indicated that the pipette tip is <100 nm from the bottom, within a TIRF illumination layer. The stationary calcium distributions were obtained at different holding potentials set to obtain a prescribed stationary calcium influx. Imaging 1D- calcium profiles was made with isotonic calcium solution in the bath and NaCl and 100 µM fluo-4 in the pipette. Ionic composition of the solutions in imaging radial calcium profiles was reversed i.e. the indicator was in the bath. In the outside-out patches the calcium patterns were similar (*n* = 6, data not shown).

### Stochastic simulations

To test theoretical predictions I simulated calcium diffusion in the presence of buffer in one- and two dimensions. The parameters of the model were *D*_*Ca*_ = 600 µm^2^/s, *D*_*fluo-4*_=300 µm^2^/s and the on-rate-constant for calcium binding *k*_*on*_ = 2.10^8^ M^−1^s^−1^. In 1D-simulations (Fig. 2A, B), a 1 μm-linear compartment was divided into 1000 cells, each contained 6 buffer molecules. This corresponds to 0.1 mM buffer, the concentration used in the experiments. The time step was 10 ns. Mean calcium levels at *x = 0* were set either to 0.2 and 2 mM. Calcium, fluo-4 and bound calcium molecules diffused, jumped into either direction chosen randomly. When calcium and free buffer molecule were in the same cell, a Ca-buffer particle was formed. Consideration of calcium unbinding did not modify the results of simulation; in line with the theoretical estimates (see Introduction). Before starting simulation, randomly set buffer molecules equilibrated for 1 ms and then calcium ‘influx’ was switched on. Time-dependent calcium patterns are plotted in Eq. 2 as kymographs, together with mean concentration profiles were obtained as averages of 1000 frames taken in the beginning and the end of the run. Two-dimensional radial diffusion was simulated in a square (Fig. 3B). Free buffer molecules at mean concentration 0.1 mM were first equilibrated for 1 ms and then calcium influx was switched on. Other parameters were the same as in the case of 1D- diffusion.

## APPENDIX Radial steady state calcium profiles

The spread of calcium from single channels into the cytoplasm is described by radial diffusion from the point source into infinite medium (Neher, 1986; Mironov, 1990; Stern, 1992). Similar to Eq. (12), the ODE for the steady state calcium profiles can be presented as

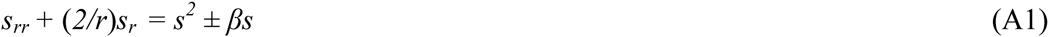
where *r* is the radial coordinate and the normalized space and concentration variables are defined by Eq. (10) in the main text. This ODE differs from 1D-case by the term in the left-hand side that contains the first spatial derivative. Substitution *s = u/r* eliminates it and gives

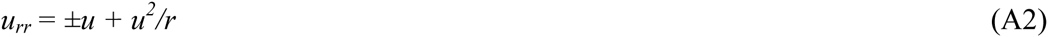

For the constant flux of calcium through the channel, the boundary condition is

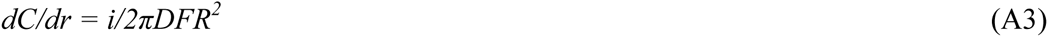
where *R* is the exit radius, *i* is single channel current, *D* is the diffusion coefficient and *F* is the Faraday constant. The substitution *C = u/r* transforms (A3) into

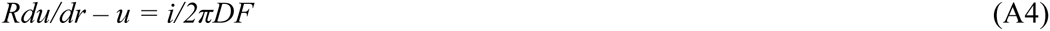

The first term in the left-hand side is small and can be neglected that defines

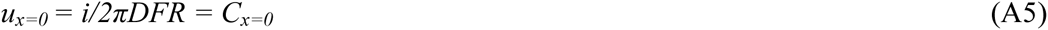

Remarkably, the condition (A5) is identical to that used in 1D-case, Eq. (8). (A2) would have the solutions identical to 1D-case, were it not the square term *u*^2^/*r*. In order to estimate its influence, I presented the right-hand side as

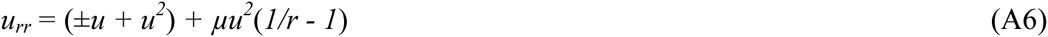
and expanded *u* = *u* + *µU*. The equation in zero order (*µ^o^*) is identical to 1D-equation (12) whose solution is given by (14) and (15). I estimated the first-order correction (*µ*^1^*U*) from the inhomogeneous ODE

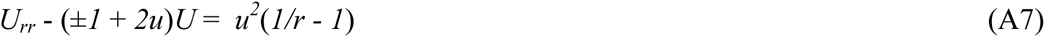
using the shooting method. Fig. A shows that for both decaying and periodic solutions the corrections in the first order are very small for all values of *A* = *C*_*x*=0_/*B*_*o*_. 1D-solutions are hence appear sufficiently accurate to obtain radial calcium concentrations after dividing them by the distance from origin.

**Fig. A.**
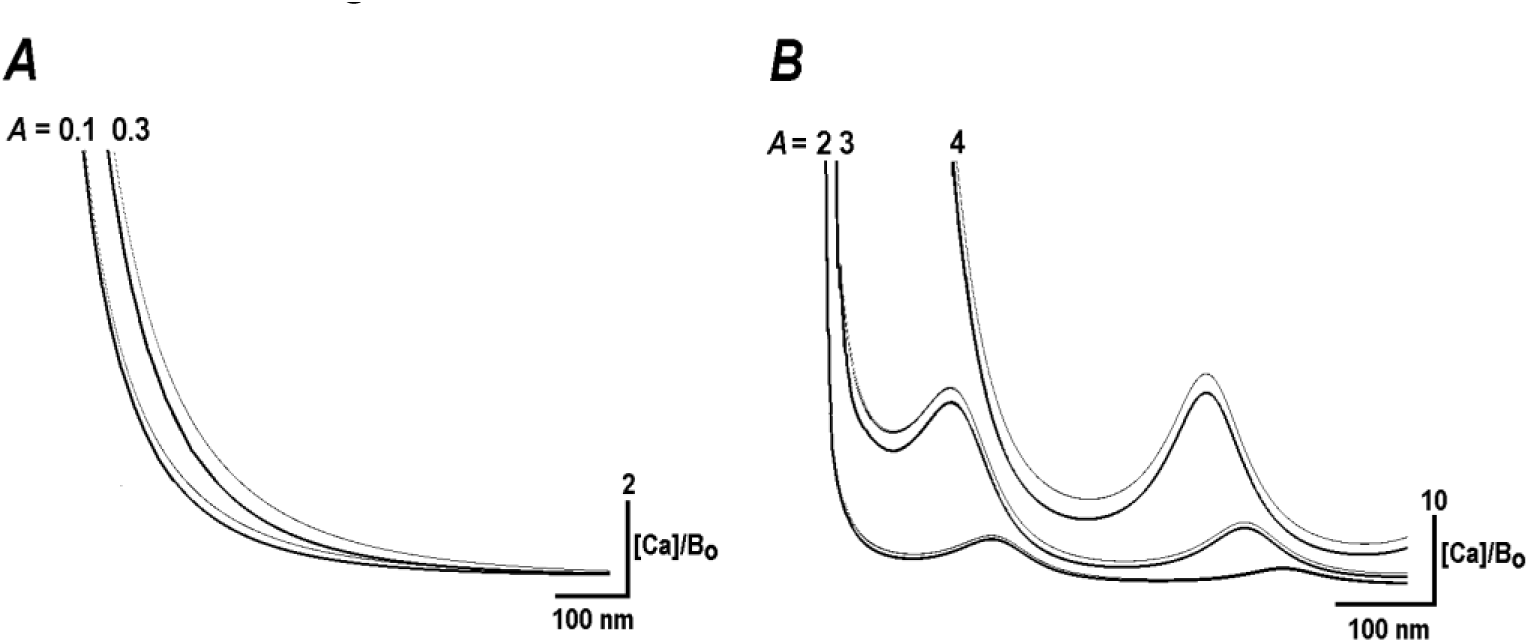
Radial calcium profiles. Thick curves present the leading (zero order) term. Dotted curves show their modification by the first order correction obtained through Eq. (A7). The calculations were made for cases when the calcium level at origin is smaller (***A***) and bigger (***B***) than the concentration of buffer (*B*_*o*_), respectively. The values *A* = *C*_*x*=0_/*B*_*o*_ are indicated near respective curves.

## References

Augustine, G. J., F. Santamaria, and K. Tanaka. 2003. Local calcium signaling in neurons. Neuron 40:331–346.

Eggermann, E., I. Bucurenciu, S. P. Goswami, and P. Jonas. 2011. Nanodomain coupling between Ca^2+^ channels and sensors of exocytosis at fast mammalian synapses. Nat. Rev. Neurosci. 13:7–21.

Beaumont, V., A. Llobet, and L. Lagnado. 2005. Expansion of calcium microdomains regulates fast exocytosis at a ribbon synapse. Proc. Natl. Acad. Sci. USA 102:10700–10705.

Demuro, A., and I. Parker. 2006. Imaging single-channel calcium microdomains. Cell Calcium 40:413–422.

Catterall WA, Perez-Reyes E, Snutch TP, Striessnig J. 2005. International Union of Pharmacology. XLVIII. Nomenclature and structure-function relationships of voltage-gated calcium channels. Pharmacol Rev. 57: 411–425.

Holmes, M. H. Introduction to perturbation methods. Springer, 1995.

Kaeser PS, Regehr WG. 2014. Molecular mechanisms for synchronous, asynchronous, and spontaneous neurotransmitter release. Annu Rev Physiol. 76: 333–363.

Koch, A. J., and H. Meinhardt. 1994. Biological pattern-formation - from basic mechanisms to complex structures, Rev. Mod. Phys. 66:1481–1523.

Korchev YE, Negulyaev YA, Edwards CR, Vodyanoy I, Lab MJ. 2000. Functional localization of single active ion channels on the surface of a living cell. Nat Cell Biol. 2000 2: 616–619.

Kostyuk PG, Mironov SL, Doroshenko PA, Ponomarev VN 1982. Surface charges on the outer surface of mollusc neurons. J Membrane Biol 70: 171–179.

Liesegang, R. E. 1896. Uber einige Eigenschaften von Gallerten. Naturwiss. Wochenschr. 11:353–362.

Mironov, S. L. 1990. Theoretical analysis of Ca^2+^ wave propagation along the surface of intracellular stores. J. Theor. Biol. 146:87–97.

Mironov SL. 1995. Plasmalemmal and intracellular Ca pumps as main determinants of slow Ca buffering in rat hippocampal neurones. Neuropharmacology. 34: 1123–1132.

Mironov SL. 2015. α-Synuclein forms non-selective cation channels and stimulates ATP-sensitive potassium channels in hippocampal neurons. J Physiol. 593: 145–159.

Mironova LA, Mironov SL. 2008. Approximate analytical time-dependent solutions to describe large-amplitude local calcium transients in the presence of buffers. Biophys J. 2008 94: 349–258.

Neher, E. 1986. Concentration profiles of intracellular Ca^2+^ in the presence of diffusible chelator. Exp. Brain Res. 14:80–96.

Polyanin AD, Zaitsev VF. (1995). Handbook of Exact Solutions for Ordinary Differential Equations. CRC Press.

Shuai, J., and I. Parker. 2005. Optical single-channel recording by imaging Ca^2+^ flux through individual ion channels: theoretical considerations and limits to resolution. Cell Calcium 37:283–299.

Smith PD, Liesegang GW, Berger RL, Czerlinski G, Podolsky RJ. 1984. A stopped-flow investigation of calcium ion binding by ethylene glycol bis(beta-aminoethyl ether)-N,N’-tetraacetic acid. Anal Biochem. 143: 188–195.

Stern, M. D. 1992. Buffering of calcium in the vicinity of a channel pore. Cell Calcium 13:183–192.

Tadross MR, Tsien RW, Yue DT. 2013. Ca channel nanodomains boost local Ca amplitude. Proc Natl Acad Sci U S A. 110: 15794–15799.

Tay LH, Dick IE, Yang W, Mank M, Griesbeck O, Yue DT. 2012. Nanodomain Ca²⁺ of Ca²⁺ channels detected by a tethered genetically encoded Ca²⁺ sensor. Nat Commun. 3: 778.

Vanag, V. K., and I. R. Epstein. 2007. Localized patterns in reaction-diffusion systems. Chaos 17:037110.

Wang, S. Q., C. Wie, G. Zhao, D. X. Brochet, J. Shen, L. S. Song, W. Wang, D. Yang, and H. Cheng. 2004. Imaging microdomain calcium in muscle cells. Circ. Res. 94:1011–1022.

